# Evolutionary rate correlations reveal long-term co-evolutionary interactions in *Drosophila melanogaster*

**DOI:** 10.64898/2026.05.21.726714

**Authors:** Andrius J. Dagilis, Selina Lee, Bonnie DiAngelis, Daniel R. Matute

## Abstract

Co-evolution between genes can occur for a variety of reasons, including co-expression of genes, epistatic interactions between them, physical interactions of gene products and many others. Co-evolutionary partners of a gene are therefore of great interest in identifying potential factors that contribute to any phenotype of interest. State-of-the-art approaches to detect these interactions use correlations of evolutionary rates across a broader phylogeny, and so by necessity identify interactions only among genes that are present across long evolutionary time periods. This makes the methods unwieldy when interest lies in a single focal organism in which the genes of interest may have evolved in the recent evolutionary past. Here, we present a new approach to calculating evolutionary rate correlations which focuses on extracting maximum coverage for a single focal species, while retaining signals of co-evolution across large clades. We show how this approach is able to identify potential interactions even in highly studied species and highly studied genes, with a focus on the *D. melanogaster* sex-determiner, *Sxl*, using data from 72 species of Dipterans.

## Introduction

The evolutionary fate of a mutation is rarely determined solely by its direct effects on fitness – often, the effect is modulated by the genetic background (Otwinowski, McCandlish et al. 2018, Johnson, Reddy et al. 2023). This is true for alleles contributing to a trait under stabilizing selection, where the fitness effect of the allele depends on its genetic background and whether it brings the phenotype closer or further from the optimum (Fisher 1958), it is true for alleles in co-expressed genes (Mähler, Wang et al. 2017a), and it is true for any case where there are epistatic interactions between different alleles (Kirkpatrick, Johnson et al. 2002, Barton 2017). In long evolutionary timespans, certain pairs of genes are more likely to show patterns of co-evolution (in this paper being defined specifically as correlated rates and topologies of evolution) than others purely due to the likelihood they are involved in similar pathways, their protein products interacting, or their gene products being co-regulated. Thus, identifying co-evolving pairs of genes can shed light onto what pairs of genes have, in the long run, been part of the same broader biological pathways.

Evolutionary Rate Correlations (ERC) are the most popular approach to identifying such co-evolutionary interactions (Clark, Alani et al. 2012, Little, Jordan, Meyer et al. 2025, Little, Jordan, Chikina et al. 2024, Hopson, Omelianczyk et al. 2026, Forsythe, Gatts et al. 2025). The general approach is to first build gene trees at each gene with orthologous sequences across the phylogeny. The gene trees are constrained to the species tree topology, and the branch lengths of the resulting trees are correlated after relativizing them to evolutionary rates using the total species tree (Clark, Alani et al. 2012). The primary issues with the method are that it: 1) only works for orthologous sequences 2) forces all trees into the same topology and 3) the combinatorial nature of the number of pairs being compared has meant the method is generally slow. The last few years have seen a few new developments in these methods, allowing them to deal with paralogs (Forsythe, Gatts et al. 2025) as well as increasing the speed and accuracy with which ERC values can be computed (Little, Jordan, Meyer et al. 2025). However, all current methods suffer from several core issues. First, because the values are calculated for a taxon rather than a species (e.g. fungi (Steenwyk, Phillips et al. 2022), plants (Forsythe, Gatts et al. 2025), *Plasmodium (Hopson, Omelianczyk et al. 2026)*), the number of genes for which ERC values are calculated naturally declines as the total number of species included in the study increases (Forsythe, Gatts et al. 2025). This is because genes need to be homologous across the whole group - with more taxa the probability that genes are missing or duplicated from some taxa increases. Even when allowing for paralogous sequences, ERC-net showed a limitation on the number of genes included versus the number of species used to calculate correlations (Forsythe, Gatts et al. 2025). The most likely genes to be excluded from ERC analyses are therefore also those that have evolved in recent evolutionary timespans, as they will have few representatives across the group, and these may be important drivers of adaptation (Zhang, W., Landback et al. 2015). Second, the fixed species tree assumption in ERC 2.0 means that genes that have evolved under an alternative topology (due to horizontal gene flow/introgression or ILS) are forced into an incorrect tree topology. When two trees are forced into the same wrong tree topology, the resulting gene trees can show rate correlations when none existed initially. This issue may be of rare concern in some systems, but evidence points increasingly to introgression being common in many systems (Dagilis, Peede et al. 2022, Suvorov, Kim et al. 2022, Edelman, Mallet 2021), . Finally, current approaches identify “significant” interactions through an outlier approach. While this allows for clear enrichment of pathways with enhanced signals of co-evolution and has proven its utility in, for instance, predicting new members of pathways (Little, Jordan, Meyer et al. 2025), they don’t allow researchers to ask what proportion of gene pairs are co-evolving.

In this paper, we propose a modified ERC approach which maximizes power for an individual species, while allowing for variation of gene tree topology and proposing a null distribution of expected signals of co-evolution. In practice, most researchers are likely to be interested in the co-evolutionary network of their system rather than the clade at large. By using a focal species, we are able to maximize the number of genes for which ERC values can be calculated by identifying reciprocal best hits with all other species in a dataset. We further develop an approach that relaxes the assumption of a fixed species tree topology for all genes, allowing each gene to have a different tree topology. While this comes at a performance cost in comparison to other approaches, it allows for ERC values to be calculated for any researcher’s species of interest. To offset this performance cost, we implemented the pipeline in Julia (Bezanson, Karpinski et al. 2012), a programming language with significant performance gains over existing implementations of ERC in Python or R. Finally, we propose two ways to create null distributions for ERC values, either as a negative null under no co-evolution, or simulations with increasing degrees of co-evolution among genes. The pipeline described here is not meant to replace either of the other available approaches - indeed, we think many of the ideas here are easily transferable to both the computationally faster ERC 2.0, and to ERC-net’s better dealing with paralogous sequences (see Table 1). However, the pipeline presented here maximizes the amount of information one can obtain for a focal species, allowing us to investigate the co-evolutionary network within a particular organism of interest, rather than across the whole clade.

**Table 1:** Comparison of existing ERC approaches and this study. Existing ERC software, like ERC 2.0 and ERC-net have distinct advantages over our study in speed and ability to deal with paralogs, respectively. However, many of the important elements of the methodology we propose here (indicated by asterisks) are transferable to the other methods as well.

| Approach | Speed | Paralogs | Focal species information | ILS/<br>gene flow | # genes | null model |
| --- | --- | --- | --- | --- | --- | --- |
| ERC 2.0 | ✓ | X | X* | X | X* | X* |
| ERC-net | ≈ | ✓ | X* | ≈ | ≈* | X* |
| This study | ≈ | ≈ | ✓ | ✓ | ✓ | ✓ |

We demonstrate the utility of this method in *Drosophila melanogaster,* a model species for which many complementary approaches have identified interacting pairs of genes. We show that we can look for co-evolutionary interactions that date back across all of Diptera (72 species spanning the whole order) in 12,511 genes. We first show that around 30% of interactions show more correlations than expected by chance alone. We then identify 1.1 million interactions (1.8%) with signals consistent with strong co-evolution. These interactions cluster into six large communities of genes, which are each enriched for different sets of biological functions, expression timings and gene product cellular localization. We demonstrate how these co-evolutionary interactions represent a separate axis to investigate genetic interactions beyond what has been previously identified in protein interaction, co-expression and other such networks. Finally, we highlight the utility of this approach by focusing on the co-evolutionary partners of *Sxl*, the sex determining gene in many dipteran species which is not a BUSCO gene and therefore would not have been analyzable with previous approaches.

## Methods

We developed a full pipeline to go from a focal species proteome to co-evolutionary interaction inference, here demonstrated for *D. melanogaster*. We describe each of the steps as follows.

### Data acquisition

The last decade has seen an explosion of available reference genomes for a variety of organisms. To maximize data quality while minimizing computational time, we focused on Dipteran chromosome scale reference genomes with RefSeq annotations. In principle, one could use whole reference genomes for the approach outlined, but we focus on annotated genomes for simplicity (and to have a “gold-standard” of protein coding sequence). We only use data from organisms with chromosome level assemblies and vetted protein annotations, but were still able to obtain data for 72 Dipteran and one outgroup (Ctenocephalides felis) species. Data was downloaded using ncbi-dataset software (O’leary, Cox et al. 2024), with the call “ncbi dataset download -taxon Diptera”. Protein sequences were then trimmed to primary transcripts using the OrthoFinder2 primary_transcript.py script (Emms, Kelly 2018). Table S1 lists the accession numbers of the data included in the analyses.

### Building gene trees

For each protein in D. melanogaster, we first performed a reciprocal diamond blast against all other species in the dataset. Reciprocal best hits were considered orthologs. Any gene with at least 5 total sequences (D. melanogaster + 4 putative orthologs) was retained for further analysis. Sequences were aligned using mafft version 7.1 (Katoh, Standley 2013). For each alignment, we built an individual gene tree using IQ-TREE2 (Minh, Schmidt et al. 2020). Trees were unconstrained to any topology, and ModelFinder (Kalyaanamoorthy, Minh et al. 2017) was used to identify the best molecular model for each sequence. The set of resulting gene trees was then used to generate a consensus species tree with ASTRAL-IV (Zhang, C., Nielsen et al. 2025), specifying *C. felis* as the outgroup. Finally, we re-root the trees using *D. melanogaster* as the outgroup - this guarantees that all trees share a common root, as it is the only species guaranteed to be present across the dataset. While this modifies tree topologies, the ERC calculations rely on branch lengths and not topologies, and re-rooting does not modify these branch lengths.

### Evolutionary Rate Correlations

We calculate a slightly modified version of ERC compared to prior approaches. Because we do not force gene trees to conform to the species tree topology, we first identify corresponding branches between each pair of gene trees and the species tree. We do this by first rooting all trees by the only node guaranteed to appear in all trees - the focal species. Because our goal is to compare branch lengths, inaccuracies in the inferred topology are not problematic as long as corresponding branches are identified consistently. We then drop any tips not shared among the two gene trees and label each branch based on its descendent tips. Branches are considered equivalent if they lead to the same set of descendants (Figure 1A). Branches need to be shared not only between the two trees, but also with the species tree, meaning we discard evolutionary rates that are not concordant with the species tree topology. Analysis for the pair is stopped if fewer than four branches are shared.

**Figure 1:**
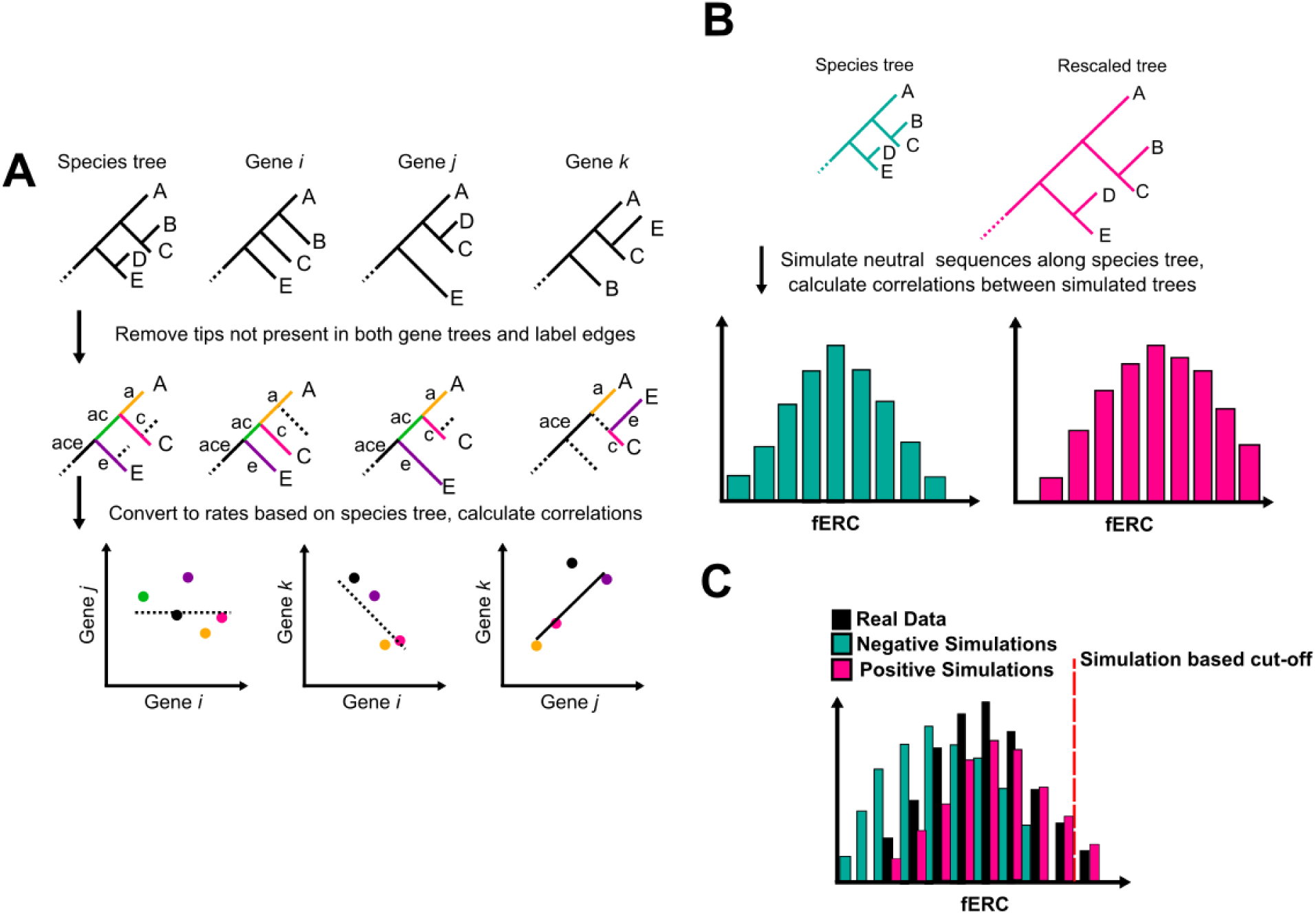
Schematic explanation of the approach in the present paper. **A)** Unlike previous approaches, we allow for tree topology to vary among individual genes. In order to calculate evolutionary rate correlations, we identify sets of branches present in both the species tree and the pair of gene trees being compared. For simplicity, we here show the branches shared by all four gene trees, with each branch represented by a different color. The branch lengths are then divided by the species tree lengths, and correlated. These correlations are then Fisher transformed to obtain the fERC values. **B)** To create both positive and negative null expectations of fERC values, we first randomly permute all branch lengths in the species tree by returning an exponentially distributed random variable with the rate parameter set to the branch length. Because these permutations are independent, the resulting fERC values between randomized trees represent a zero co-evolutionary interaction background. We then rescale the species tree branch lengths and run coalescent simulations along the resulting tree. Because of long coalescent time periods, the resulting simulations give trees with both highly similar topology and branch lengths, representing a positive null for pairs of genes evolving similarly due to shared genetic background selection, mutation or recombination rates. **C)** The actual fERC values are then compared to a simulation informed cut-off (95th percentile in this paper) to determine significantly evolving pairs of genes.

We next calculate ERC as in previous approaches – branch lengths in each gene tree are divided by the branch length in the species tree to account for lineage-wide evolutionary rate variation. We discarded branches with values over five (representing extremely rapid evolutionary periods), as these can generate false positive correlations. Finally, we compute a Pearson correlation between the two genes’ evolutionary rates. Following the precedent of ERC 2.0, we calculate a Fisher’s Transform of the correlation as in ERC2.0 (Little, Jordan, Meyer et al. 2025), allowing more direct comparisons between pairs of genes with different numbers of shared branches. We refer to these transformed values as fERC and use them for all downstream analyses. The code for fERC calculations was written in Julia and is available at: https://github.com/adagilis/CoEvolution

### Null fERC calculation

While each pair of genes in the above analysis will include both a correlation and a p-value for a correlation test, the combinatorial nature of the analysis results in many millions of pairs of genes being examined, and thus an appropriate null model is desired to avoid false positives resulting from the large number of comparisons. We generate two sets of expected fERC values – one under the assumption of independent evolution between the pair of genes, the second assuming evolutionary rates are highly similar.

#### Negative null distribution under no correlation

For the first approach, we generate a permuted set of branch lengths from the species tree, assuming each branch length is distributed with an exponential distribution, with the current branch length as the rate parameter. This creates a set of independent branch rates in which any changes are not correlated. A true null including variation in tree topology would allow for topologies to vary when branch lengths are short as well. In practice, we found that this resulted in very few shared branches between most pairs of trees, which returns missing values rather than correlations of 0. This biases the distribution towards pairs of genes that happen, by chance, to share topology, and therefore often branch lengths. Thus, for a null distribution we keep the topology fixed to the species tree but allow branch lengths to fluctuate independently. We permute 1,000 such trees and calculate the resulting fERC values for 499,500 pairs of trees. We use the resulting distribution as a first-pass filter: any pair of genes in our fERC data with values below the 95 percentile of the null distribution are discarded.

#### Positive null distribution with correlated evolutionary rates

Second, we perform coalescent simulations along the species tree with branch lengths scaled 10x. We use the PhyloCoalSimulations package (Fogg, Allman et al. 2023) to run coalescent simulations. The package simulates a set of trees given an input tree with branch lengths scaled to coalescent units. We scale the branch lengths 10-fold to cause much longer coalescent times, and therefore, higher degree of correlation between gene trees than expected by random chance. We simulate 1000 such trees and calculate fERC values for all pairwise combinations of the resulting set of trees (resulting in a positive distribution of 499,500 pairs of trees).

#### Significance testing

We finally assess significance for each gene pair using a combination of its fERC value compared to the positive simulation as well as the Benjamini-Yekutieli adjusted p-values (Benjamini, Yekutieli 2001) from the Pearson correlation. Gene pairs with fERC scores above 95% of positive simulations, and an adjusted p-value < 0.05 are considered to be significantly interacting.

### Network analyses

For pairs of significantly interacting genes, we construct an undirected network with each gene representing a node, while edges represent significant interactions. We use Leiden clustering (Traag, Waltman et al. 2019) to find communities of genes. Leiden clustering identifies communities by maximizing the modularity (relative number of edges within vs between communities). Clustering is performed using the igraph library in R (Csardi 2013), and the cluster_leiden function and 10,000 iterations.

Each gene was annotated for its community membership and degree (number of interactions). We further calculated the average fERC value for each gene with all other genes tested.

#### GO enrichment

We look for GO-term enrichment within our network in several complementary ways.

For each community identified in our network analysis, we tested for Gene Ontology (GO) term enrichment using the gprofiler2 package in R (Kolberg, Raudvere et al. 2020), with “dmelanogaster” as the organism, and a custom gene background set to either all genes included in our pipeline (12k genes), or, for a more conservative outlook, genes in one of the major communities (8k genes). Outputs for these analyses (Tables S4,7) were examined for the highlighted GO categories, which may be the drivers of other GO category enrichment (g:profiler highlighting description).

When examining the interactions of a single gene, GO enrichment using solely significant interactions may be inaccurate, because some GO terms may simply have higher expected degrees of co-evolution. For instance, genes involved in protein transport are likely to co-evolve with many other genes. Seeing enrichment for “protein transport” for the interaction partners of a single gene is therefore not an indicator of meaningful enrichment for that gene. We therefore build a null-expectation of fERC for each GO-term. We first downloaded the latest version of GO annotations for *D. melanogaster* (Thurmond, Goodman et al. 2019). For each GO-term, we calculate the mean fERC value of each gene with that GO-annotation using a custom script. Then, to look for enrichment for any gene’s interactions, we compare its observed fERC values with all genes in each GO term with the expected fERC values given the mean fERC of those genes. We perform a Mann-Whitney U test to look for GO-terms for which the gene has fERC values that are distinct from expectation, and perform Benjamini-Hochberg FDR control at a 0.05 level to look for terms that are significant. Custom scripts to perform this analysis were written in Julia.

## Results

### fERC values in *D. melanogaster*

Out of 13,962 protein coding sequences listed in the *dmel6* reference genome annotation, we found 12,511 with at least 4 reciprocal-best-hit orthologs among our 72 species of *Dipterans*. We built gene trees for each ortholog group and calculated all 78,256,305 pairs of fERC scores on the resulting gene trees. Of these, 71 million had sufficient shared evolutionary branches (>4) to calculate a correlation score. Around 30% of the resulting interactions exceeded our negative null. We found an overall low density of interactions (∼1.5%) (Figure 2A) exceeding our positive null distribution of genes evolving under similar evolutionary rates, yet this still represented nearly 1.2 million pairs of genes with significant co-evolutionary interactions.

**Figure 2:**
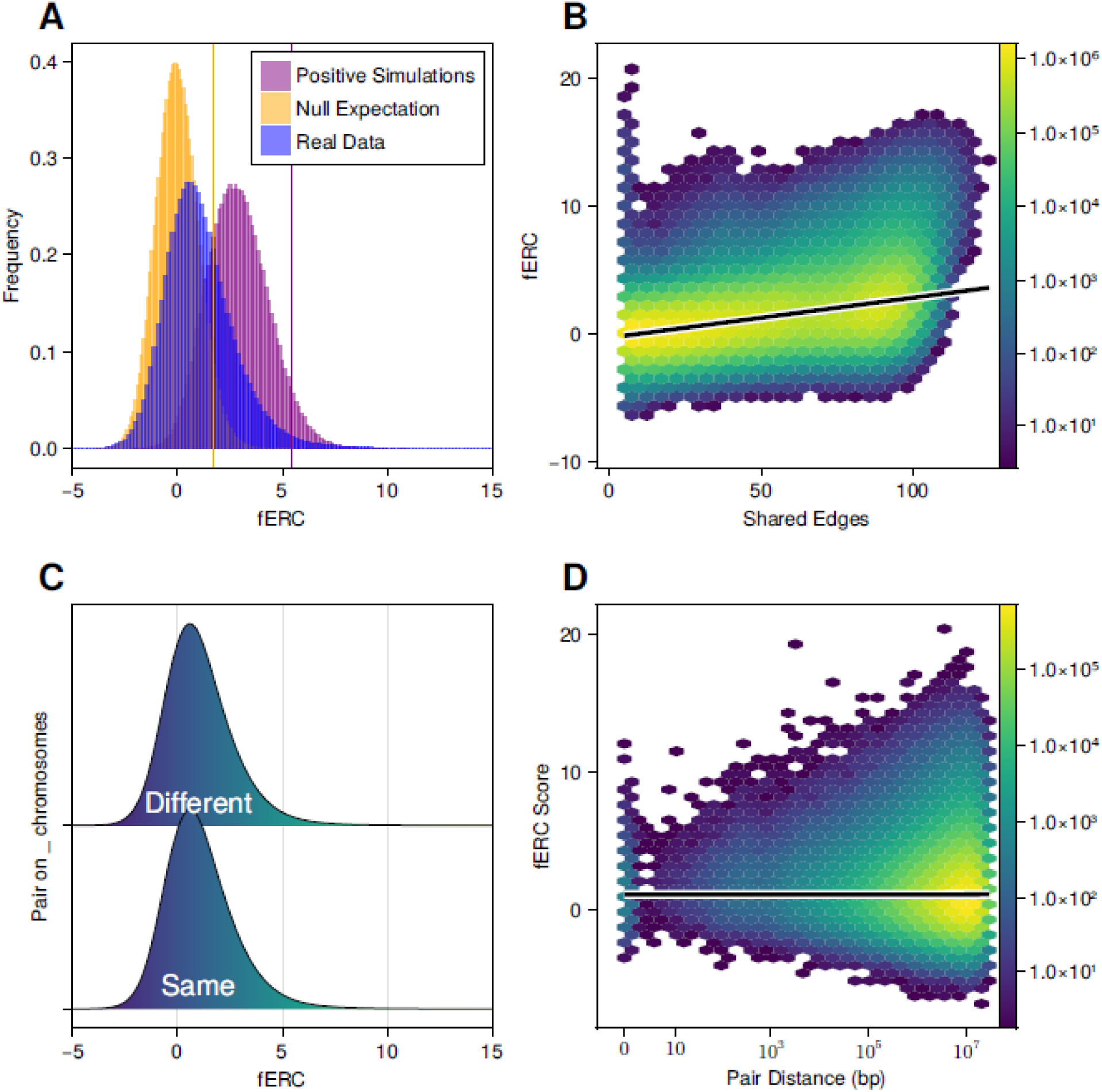
The distribution of fERC scores in the *D. melanogaster* data. **A)** The overall distribution of fERC scores lies intermediate to the negative and positive null simulations - while many pairs of trees show higher correlations than under a model of no correlation, they show lower signals than may be expected by chance. **B)** The distribution of fERC scores given the number of shared edges in gene trees (colors represent frequency of pairs of genes in each bin, line is the best GLM fit). Overall, pairs of genes which share more edges in their gene trees show slightly larger values of fERC. However, it’s worth noting that this is likely due to a true shared history leading to more shared edges, rather than a systemic issue with more shared edges driving larger fERC values (see Figure S1). **C)** The distribution of co-evolutionary interactions for gene pairs located on the same vs different chromosomes. Gene pairs on the same chromosome have marginally larger fERC scores (mean 1.09 vs 1.06, Mann-Whitney U test p-value <10^−10^). **D)** For pairs of genes on the same Muller element in *D. melanogaster*, fERC scores generally increase with increasing genetic distance, although the effect size is minimal (0.09 fERC/Mb). Colors indicate frequency of pairs of genes in each bin; the line is the best GLM fit.

As a validation step, we examined the genes with the largest average signal of co-evolution (Tables S2,S3). The top genes fall into functionally coherent categories (Table S3). Multiple heat shock proteins fall in the category of most highly co-evolving, and heat shock proteins are likely to have to interact with many other proteins in correcting/preventing mis-folding during a heat response. In yeast, individual heat shock proteins physically interact with up to 10% of all genes (Taipale, Jarosz et al. 2010). Other genes include chromatin remodelers, ATP binding cassettes and glycogen synthase.

We find that gene pairs with greater shared evolutionary history–as reflected by a larger number of common tree branches – show higher fERC scores overall (Figure 2B). This pattern can arise for two reasons. First, a larger number of shared branches might also lead to an increase in statistical power to detect correlations thus raising the likelihood of detecting concordant rate variation among gene genealogies. On the other hand, if coevolving genes tend to share similar tree topologies, our decision to allow topology variation may preferentially retain pairs with greater topological concordance, thus inflating both the number of shared branches and fERC values. To distinguish between these explanations, we estimated the correlation between the number of shared orthologs (reciprocal best hits) and the number of shared edges in the tree. The ratio of retained edges is positively correlated with the number of shared orthologs (Figure S1), suggesting that increases in fERC are more strongly associated with topological concordance than just with the presence of orthologs across taxa. Overall these results indicate that fERC values are higher when there is more shared history among the pair of genes, rather than an indicator of increased power to detect co-evolution when more information is present.

Next, we evaluated if coevolutionary networks are more likely to colocalize to the same chromosomes. Overall, we find no strong indication that genes on the same chromosome, or even in closer proximity, are more likely to interact (Mann-Whitney U-test: p-value <10^−10^, point estimate = 0.025 ; p-value of fERC ∼ distance(bp) < 10^−30^, R^2^=10^−5^, slope = 9.09^−10^ fERC/bp). This is somewhat unexpected, as one mechanism generating correlated evolutionary rates is genomic proximity (and therefore shared local mutation/recombination rates). However, given that LD-decay occurs very rapidly in *D. melanogaster* (r^2^<0.5 within 500 base pairs (Ometto, Glinka et al. 2005)), it is likely that nearby regions evolve rather independently. Further, by taking a conservative approach and looking only at pairs of genes that exceed a 95% cutoff of a positive simulation, we aimed to minimize potentially weak signals arising from shared genomic neighborhoods.

We do find that interactions within chromosome 4 (the dot chromosome) are enriched (Supplementary Figure 2). Chromosome 4 frequently appears as an outlier in studies of *D. melanogaster* (reviewed in (Riddle, Elgin 2018)), so this may represent a true signal, although it could be driven by its small size and relatively low gene content. Additionally, we observed fewer significant interactions for mitochondrial genes. This result could also be driven by the relatively small number of such genes in our data (13), or due to differences in mito-nuclear dynamics across species pairs that may generate heterogeneous coevolutionary signals. Multiple communities of interacting genes seem to be enriched for mitochondrial function genes (Table S4).

In general, we do not detect any broad genomic regions with significant enrichment for elevated fERC values (Figure 3, left panel). A notable exception are centromeric regions, which show slightly elevated fERC values with themselves and other centromeres (Figure 3, upper triangle). This pattern is consistent with the expectation that centromeric regions experience coordinated increases in evolutionary rates during episodes of centromere turnover, which can affect multiple centromeres simultaneously. In *Drosophila*, centromere turnover has likely contributed to karyotype evolution (Bracewell, Chatla et al. 2019, Kyriacou, Heun 2023), supporting the possibility of shared evolutionary rates among centromere adjacent genes. Despite this, genes within centromeric regions do not show more significant interactions with each other than with genes in other regions (Figure 3, lower triangle), nor do they show an increased number of significant interactions on average (Figure 3, lower panel). Put together, these results suggest genomic proximity does not drive evolutionary rate correlations in this system.

**Figure 3:**
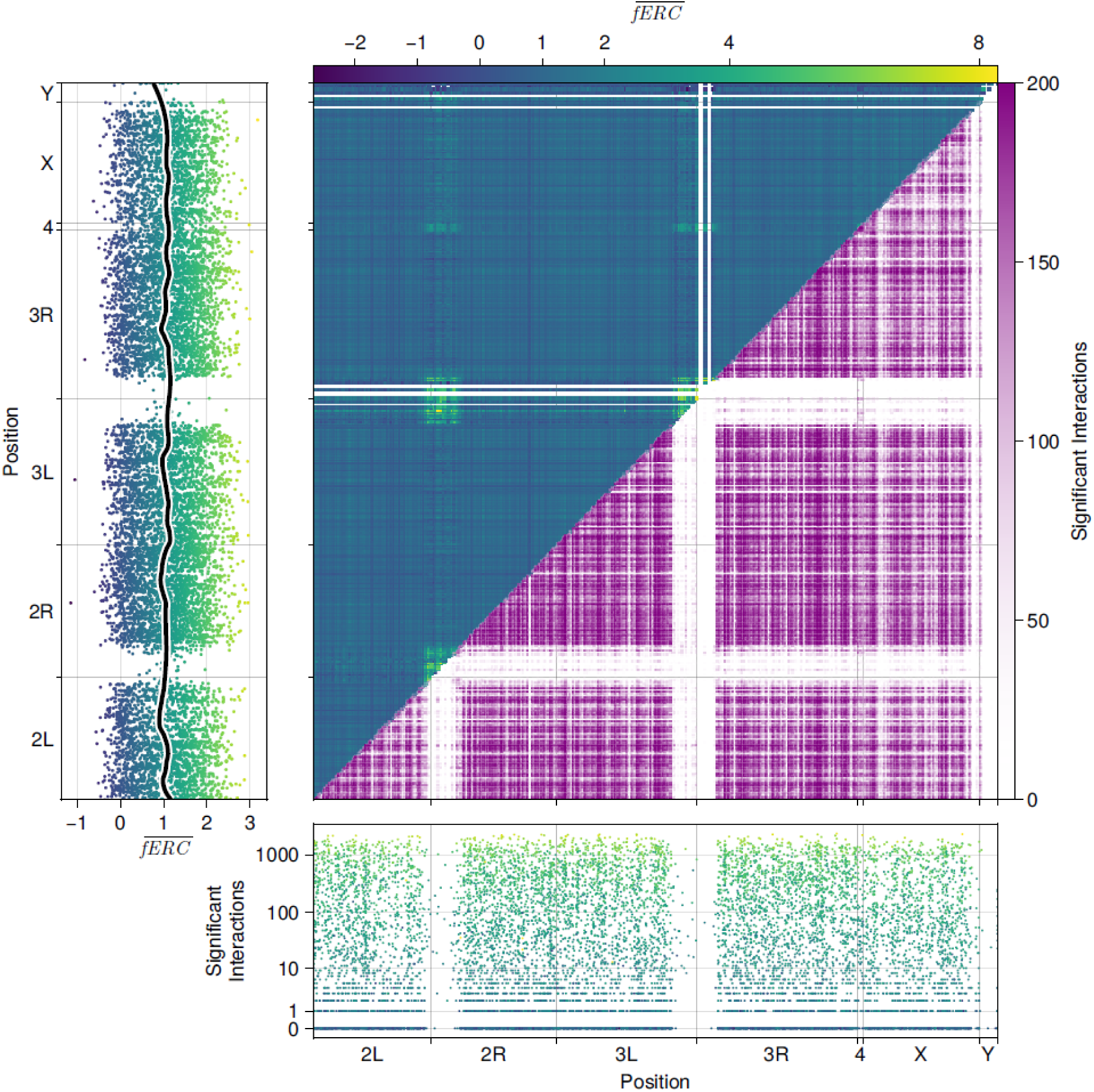
Overall genomic patterns of co-evolution. Average fERC values by gene (left), across different genomic regions (middle panel, top triangle), as well as significant interactions per gene (bottom) and between different genomic locations (middle panel, bottom triangle) show little pattern to concentration of co-evolutionary interactions across the genome. Pairwise correlations are slightly elevated among centromeric regions (middle panel, top triangle), but centromeric genes have few significant interactions (bottom panel). For a summary of interactions by chromosome see Figure S2.

### The co-evolutionary network of *D. melanogaster*

Next, we determined what proportion of interactions exceeds a negative null expectation of fERC scores. In this null model, any correlations between two gene trees are entirely spurious. We find that 29.8% of interactions exceed the 95% cut-off generated by this null. We next simulate a set of trees under a coalescent model. Each gene tree is simulated under the species tree, with coalescent rates scaled down ten-fold. This generates gene trees with, on average, higher correlations in branch lengths as the longer coalescent times lead to less variation in branch lengths and topology.

Following our significance testing, we retain 1,179,974 interactions among 8859 genes. The network of these interactions largely forms a single connected component (Figure S3), alongside a smaller set of disconnected gene pairs. The overall network structure is consistent with expectations for biological systems - the degree distribution and clustering coefficient more closely resemble those of a small-world network than a random network, in which the degree distribution (number of nodes with significant partners) follows a Power Law (Figure S4). The main component (set of connected genes) of 8754 genes clusters into five communities with more than 50 members (Table 2), identified as groups with more interactions within each group than to other such groups. These communities seem to represent sets of sequences with different evolutionary histories – we generated consensus phylogenies using ASTRAL-IV for each community (Figure S5A) and plotted their relative evolutionary rates compared to the species tree (Figure S5B). Community 1, for instance, seems to overall match well with the species tree, while community 2 represents slowly evolving sequences that nonetheless largely follow the species tree topology.

**Table 2:**
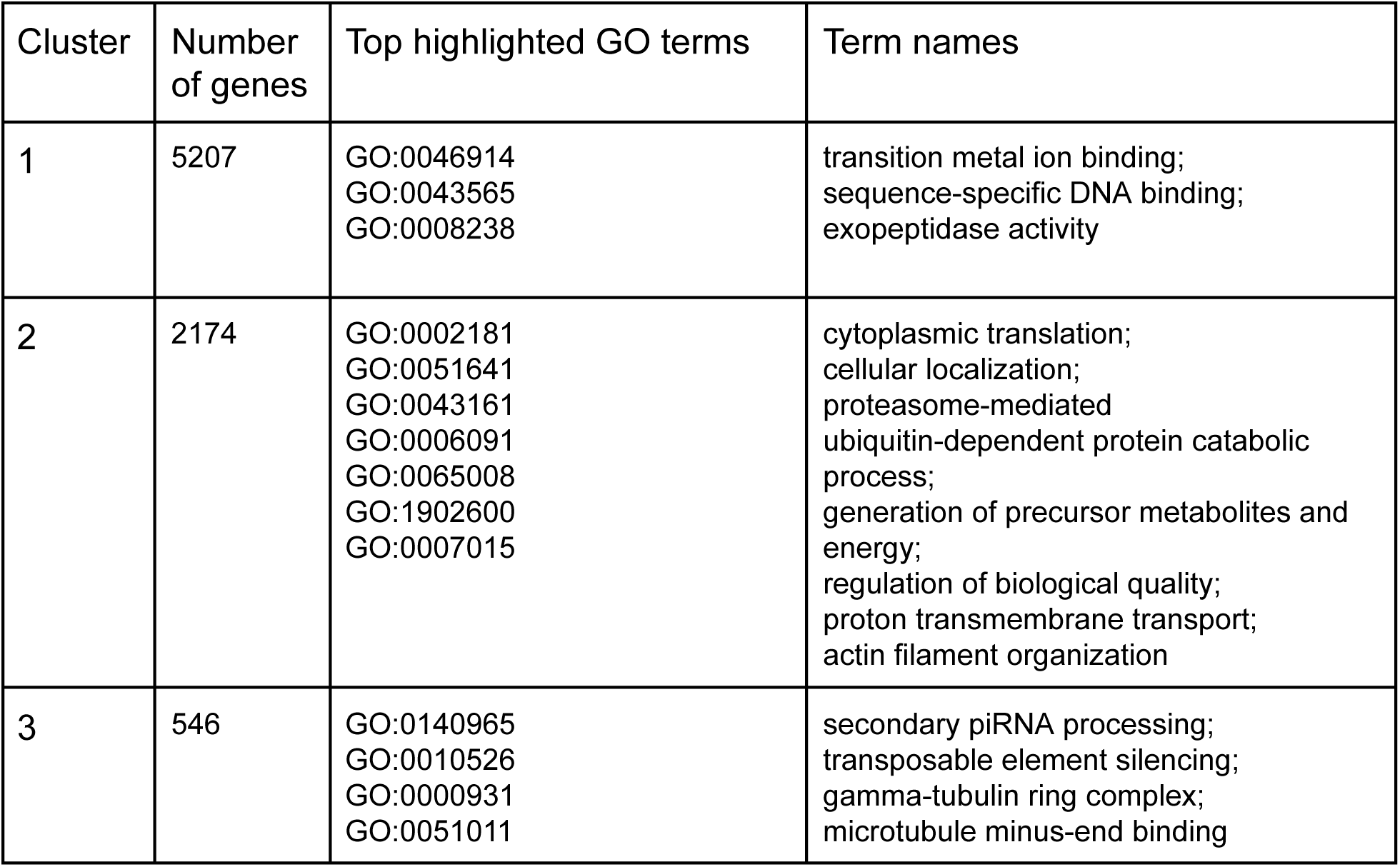
Putative drivers of GO enrichment among the five clusters. Drivers identified using *gProfiler2*’s highlight function. Communities 4 and 5 had no highlighted terms.

For each of the five communities with more than 50 members, we performed GO enrichment analysis using *gprofiler2*. All communities show significant enrichment across multiple categories, accounting for the restricted genomic background of the genes within the largest connected component. In total, we identified 293 terms enriched across the five communities. To avoid *a posteriori* interpretation, we used gprofiler’s approach of identifying driver terms for each cluster. This revealed 30 terms that capture the major functional signals (Table 2), with none in community 4 or 5. Because community 1 contains the vast majority of the co-evolving genes, it shows relatively little enrichment.

### *Sxl* co-evolutionary partners

We next demonstrate the utility of the ERC approach by focusing on a single gene of interest - the sex determiner *Sxl (Salz, Erickson 2010).* As a master regulator of sex determination, *Sxl* is expected to co-evolve with a variety of genetic partners, including those involved in sexual differentiation as well as genes under recurring sexually antagonistic selection (van Doorn, Kirkpatrick 2010, Rice 1987) Because *Sxl* is located on the *X*, two-thirds of its copies reside in females, and only one-third in males. This may also mean that *Sxl* can co-evolve with genes associated with maternal inheritance, meiotic drive of the X and similar processes tied to the sex chromosomes.

Overall, *Sxl* does not show strong signals of increased co-evolution compared to the rest of the genome (Figure 4A). *Sxl* co-evolves more strongly with other genes that show stronger signals of co-evolution (Figure 4B), with a notable exception of *RpS23*, a gene with one of the lowest average fERC scores that is positively co-evolving with *Sxl*. Five genes show significant fERC values when compared to our positive null - *DMAP1*, *Chd64, syd*, *Deaf1* and *snx3* (Figure 4C). These genes also seem to show specific co-evolution patterns with *Sxl* (large, positive fERC_Sxl_-fERC_mean_). The genes all seem to play roles in regulatory functions - *snx3* is involved in the secretion of morphogens *(Zhang, P., Wu et al. 2011)*, *Deaf1* is a crucial transcription factor of early development (Veraksa, Kennison et al. 2002), *DMAP1* is a chromatin remodeler (Goto, Fukuyama et al. 2014), *chd*64 is involved in juvenile hormone signaling (Li, Zhang et al. 2007) and *syd* is an organ size regulator (Ahmad, Vadla et al. 2021). The small number of these interaction partners makes it hard to ask whether particular GO terms are truly enriched for co-evolution with *Sxl*, so we re-analyze the data using a different approach to gain a broader picture.

**Figure 4:**
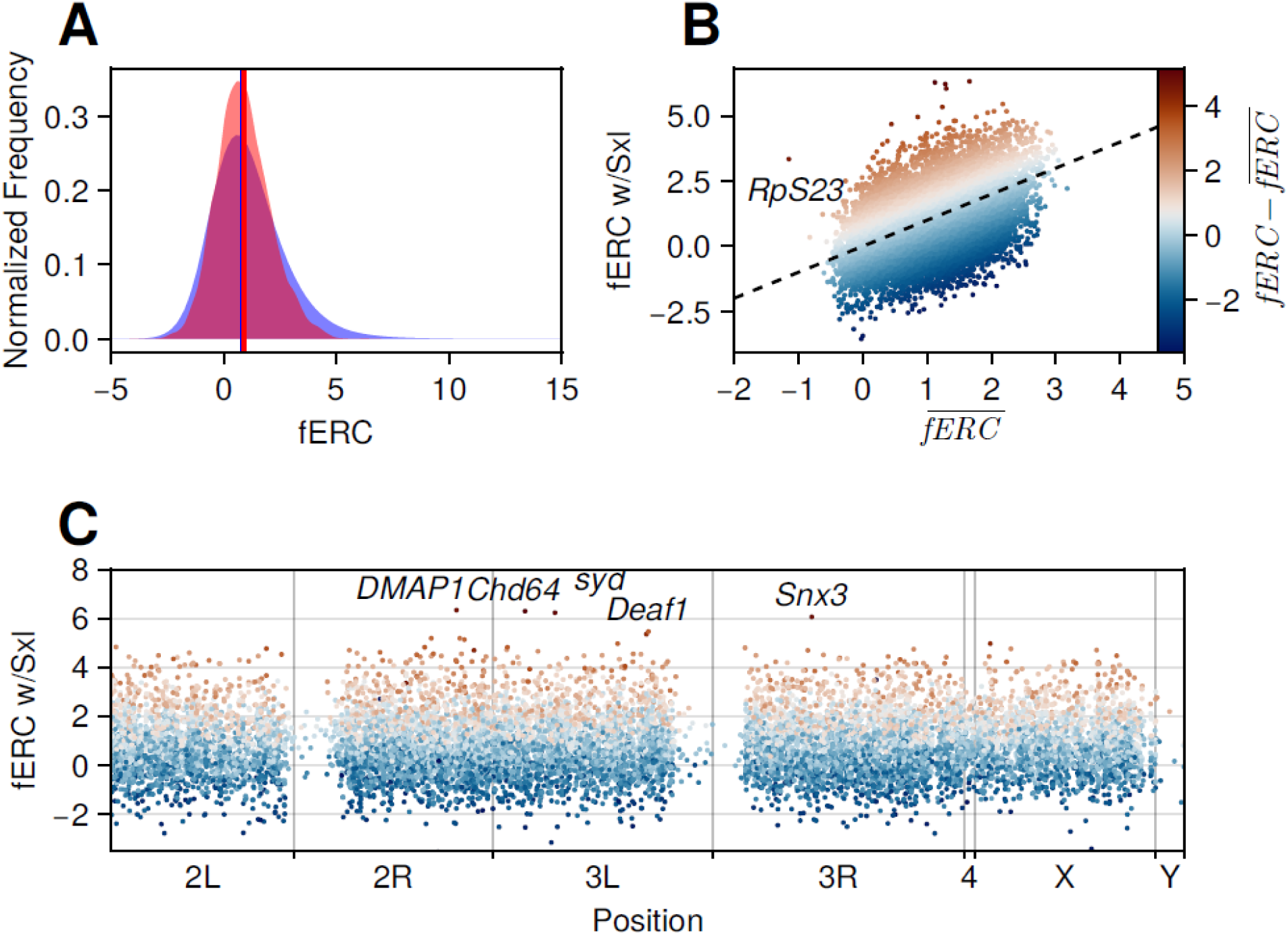
Co-evolution with *Sxl*. A) fERC values for *Sxl* (red) are distributed similarly to the genome-wide interactions (blue), with the mean interaction marginally higher than average. B) Overall, fERC with *Sxl* is correlated to the average fERC of each gene. One notable exception is *RpS23*, which on average has negative fERC with other genes, but a highly positive fERC with *Sxl*. C) Elevated fERC values with *Sxl* are found across the genome, with no clear outlier regions. The five genes with significant interactions compared to our positive simulations are annotated, and color indicates the difference of fERC with *Sxl* versus average fERC for that gene as in B.

We first take a broad view of interaction strengths involving *Sxl*. For all 12,511 genes with fERC values for *Sxl*, we test whether interactions with specific GO terms differ from background expectations. These analyses revealed only three GO categories with elevated co-evolution with Sxl: GO:0022626 (CC:cytosolic ribosome), GO:0022625 (CC:cytosolic large ribosomal subunit), GO:0002181 (BP: cytoplasmic translation). In contrast, many categories show lower-than-expected co-evolution with *Sxl* (Table S6). While some of these terms may represent genes unrelated to *Sxl* function, several patterns are noteworthy. The most negative co-evolution between *Sxl* and a GO category is GO:0046680: response to DDT. This signal is not driven by average negative fERC values in the response to DDT category, but rather by strong negative associations between *Sxl* and several of the genes in the group. Four other terms show similarly significantly negative co-evolution with *Sxl* despite positive fERC expectations: GO:01063890 - ecdysteroid 22-kinase activity, GO:0007378 - amnioserosa formation, GO:0015277-kainate selective glutamate receptor activity, and GO:0010171 - body morphogenesis (Table S6). See the discussion for what negative correlations in evolutionary rates could mean.

Finally, we examined genes with large positive interactions with Sxl. We identify 309 genes with positive, and significant fERC values exceeding the 50% threshold of our co-evolutionary simulations and tested this set for GO enrichment using gProfiler2 analysis on the resulting set of genes. We find enrichment for 94 GO terms (Table S7), including expected terms such as oogenesis, female gamete generation, and multicellular organism reproductive development.

## Discussion

In this study, we have shown how evolutionary rate correlations can be used to identify genetic interactions in focal species. Our approach maximizes the number of genes for which interactions can be detected, and couples it with a two-pronged null model to allow for the identification of interactions whose strength exceeds either neutral expectation, or expectation given a conservative null model. Our approach seems to be robust to potential drivers of correlations between genes due to their proximity in the genome, and identifies interactions that correspond to clear drivers of co-evolution (co-localization, shared genetic pathways, etc.). While this current study was performed in *D. melanogaster*, a species with rich genetic network information, the method should be easily scalable to non-model organisms, allowing for rapid identification of candidate genes in systems without pre-existing information on physical interactions, co-expression or co-regulation networks. Our work follows several recent developments in co-evolutionary network detection, and while our approach is neither as fast as ERC 2.0, nor are we able to account for paralogs as cleanly as ERC-net, we show improvements in both creating an appropriate null model and maximizing the number of genes for which interactions can be examined.

Several caveats should be considered when interpreting these results. First, to maximize signal for the focal species, we identify orthologs using a reciprocal best hit approach. Although this approach often recovers the correct orthologs, the presence of paralogs in other species could alter the inferred co-evolutionary signal. Consider a gene which co-evolves with genes in sets A and B in the focal species. If a duplication in some clade allowed each copy to subfunctionalize to interactions with B, restricting the analysis to the reciprocal best hit could make it appear that the gene co-evolves only with set A, because the signal for set B is masked. In turn, our approach may miss certain co-evolutionary interactions, although overall, it is unlikely to inflate the signal.

A second concern is that while multiple studies have found that ERC and (by extension) fERC correlate well to a variety of true interactions between genes (Little, Jordan, Meyer et al. 2025) confirming “true” co-evolution is in practice nearly impossible. It is useful to think of evolutionary rate correlations as evidence of correlated evolution, while recognizing that such correlations may arise from processes other than direct co-evolutionary interactions. This could be due to shared changes in mutation/recombination rates, although at least in our data genetic proximity in *D. melanogaster* does not impact fERC (although, note that there are few syntenic regions in Diptera, with gene order commonly being scrambled even considering closely related species (Kim, Wang et al. 2021)). Testing across a broader set of systems could verify if this pattern holds more broadly.

Third, while our approach allows us to gauge how many interactions exceed a null distribution of no co-evolution, it is difficult to gauge how many of the interactions exceeding this null are causal, and how many are a result of cross-correlations (e.g. if A co-evolves with both B and C, it is likely that we would also detect correlations between B and C). This issue is enhanced by the fairly broad range of species we examine: co-evolution anywhere along the tree can be enough to drive signals. It is unlikely that 1:3 gene pairs in *D. melanogaster* have co-evolved - rather, 1:3 pairs of genes have co-evolved for at least part of *Dipteran* evolution. It is worth making this point more broadly - while we use a focal taxon to maximize the number of genes investigated, the co-evolutionary signal is not species specific, but is determined by the taxon. The results obtained in this paper show the co-evolutionary patterns of *D. melanogaster* genes across *Diptera*, rather than identifying co-evolution specific or unique to *D. melanogaster*. The branch leading to the focal taxon represents only a single datapoint, and so the vast majority of co-evolutionary signal depends on the choice of taxon, not focal species. With these caveats out of the way, there are some notable results stemming from our analysis of *D. melanogaster* genes across Diptera.

The most surprising result of our study is the sheer number of gene pairs with fERC scores exceeding the null distribution of no evolutionary correlations. The methodology used to calculate fERC is meant to account for several potential drivers of such correlations. We calculate fERC using branch lengths supported by the species tree and both of the gene trees, and then comparing rates divided by species tree branch lengths. The resulting correlations should therefore be caused by shared changes in evolutionary rates rather than distinct evolutionary histories or genome-wide changes to evolutionary rates. Roughly a third of gene pairs have correlations that exceed a null of branch lengths drawn randomly based on the species tree. This could be interpreted as 1 in 3 pairs of genes co-evolving, but a more likely explanation is that there are many spurious cross-correlations. That is, if gene A co-evolves with both B and C, a spurious correlation may also exist between the evolutionary rates of B and C. The 1 in 3 figure therefore represents an upper boundary to the causal number of co-evolutionary interactions. Nonetheless, the large fraction of gene pairs that show correlated evolutionary rates is informative of the evolutionary process. Few genes may be evolving independently even if the number of causal interactions is smaller. Considering our previous example - if evolution at B causes elevated evolution at A, this may also drive further downstream evolution at C albeit at lower rates. Similarly, strong constraint at C may limit evolution in B due to shared interactions with A. The large fraction of correlated pairs of genes suggests that long-term evolutionary trajectories are often impacted by selection pressures at other loci. This is consistent with classical results demonstrating the genes with broader expression profiles evolve more slowly than those with narrow profiles (Mähler, Wang et al. 2017b).This may also be a part of the explanation for why observed genetic diversity rates are far lower than expected (termed Lewontin’s Paradox). One explanation for this pattern is strong background selection - purifying selection at one locus reducing nearby genetic diversity. Recent work has suggested that given recombination rates in most species, such selection is insufficient to explain the paradox (Buffalo 2021). If diversity is also constrained due to broad co-evolutionary interactions, then selection at one locus may depress diversity at loci across the genome. Further work examining how this fraction of interacting pairs of genes compares with other species in other taxa is necessary to see if this is a pattern unique to *Diptera*, or is more broadly expected. Previous fungal studies showed vastly different distributions of ERC depending on the methodology used (Steenwyk, Phillips et al. 2022, Clark, Alani et al. 2012, Little, Jordan, Meyer et al. 2025), and the single study in *Plasmodium* showed a distribution of ERC values centered firmly around 0 (Hopson, Omelianczyk et al. 2026), suggesting little to no systemic co-evolution in that system.

Looking for interactions that exceed a background of positive correlation may therefore allow for clearer identification of causal interactions. In our case, only 1.5% of interactions show fERC values exceeding simulations under a species tree with 10x longer branch lengths (and therefore highly reduced variation in evolutionary rates for individually simulated gene trees). The combinatorial nature of pairs of genes means that this 1.5% still represents over 1 million pairs of genes, spread widely throughout the entirety of the *D. melanogaster* genome with no clear enrichment of interaction hotspots. The notable exception is chromosome 4, which shows many more significant interactions within the chromosome than expected by random chance alone. If we examine the “specificity” of each interaction (calculated as the observed fERC minus the mean of both interaction partners), we find that these centromeric and chromosome 4 interactions are also highly specific (Figure S4), while interactions across the rest of the genome show few, if any, regions with either elevated or decreased signal of co-evolution. The network of significantly interacting genes shows features typical of biological interaction networks, with many genes that show overall few interactions, and a few “hub” genes. These hub genes seem to fall into categories that make sense (e.g. heat shock proteins, Table S2). Clusters of genes within this co-evolutionary network are enriched for a variety of GO-terms (Table S4), but enrichment depends in part on our choice of clustering algorithm, and assigning each gene a single “community” of interaction partners is unlikely to represent biological reality for most genes.

We can therefore look at individual gene’s fERC profiles, providing novel insights into their evolutionary history. In our case, we identify 6 outliers in the interaction profile of *Sxl*. Five genes exceed fERC values expected from positive simulations (Figure 4). These genes correspond to a wide array of important developmental processes, with plausible importance for both direct sexual function and sexual dimorphism in Dipterans, but none are core sex determination genes. *Sxl* shows little enrichment for co-evolution with any particular GO term, with its only significant terms relating to cytosolic ribosome translation. In *D. melanogaster*, *Sxl* targets both the splicing and translation of *msl-2* (Salz, Erickson 2010), the latter of which occurs in cytosolic ribosomes. Notably, *Sxl* co-evolves much more strongly than expected with *RpS23*, a component of the small ribosomal sub-unit which *Sxl* interacts with, but *Sxl* is not known to physically interact specifically with *RpS23* in *D. melanogaster* (BioGrid: rps23, Oughtred, Rust et al. 2021). The remaining genes *Sxl* co-evolves with significantly are, to the authors’ knowledge, not known to have clearly sexually dimorphic roles. *Sxl* shows lower than expected (and indeed negative) correlations with genes in the “response to DDT” GO term. DDT has sexually differentiated outcomes in *Drosophila* (Rostant, Kay et al. 2015, Rostant, Bowyer et al. 2017), raising the possibility that sex-determination pathways may have been constrained under insecticide exposure, or vice versa. Finally, GO enrichment of genes with elevated fERC values shows that the terms with highest recall (largest proportion of all genes in terms that are significantly co-evolving with *Sxl*) included genes involved in actin organization/regulation/localization as well as viral processes (Table S7). Actin plays an important role both in meiotic drive (with actin binding often determining which chromosomal copy ends up in the polar body or the egg), and may play an important role in endosymbiont regulation (including *Wolbachia* and *Rickettsia* (Newton, Savytskyy et al. 2015)). The unexpected genes and gene categories we find co-evolving with *Sxl* demonstrate that even for an extremely well studied gene in a model system, examining co-evolutionary interactions may provide novel insights into both ultimate and proximate questions.

In general, previous studies have only focused on the positive rate correlations between pairs of genes (Clark, Alani et al. 2012, Little, Jordan, Chikina et al. 2024, Little, Jordan H., Meyer et al. 2025, Hopson, Omelianczyk et al. 2026, Steenwyk, Phillips et al. 2022, Forsythe, Gatts et al. 2025). It is unclear exactly what a negative fERC value represents. Negative rate correlations suggest that for some subset of gene tree branches, one gene is evolving more rapidly than expected, while the other more slowly. One simple explanation is that the pair of genes are under some form of evolutionary constraint (evolution in one slows the other), and such interactions are surely of interest. An alternative is that spurious signals are caused due to horizontal gene transfer/common introgression. Suppose A evolves strictly under the species tree, while B undergoes HGT. B may show many shorter divergences than A between distant taxa when gene flow from a distant species has occurred, and consequently highly different branch lengths than A. By allowing tree topology to vary for each gene, we should be able to remove most false signals from such events since many branches between the resulting trees should simply not be shared. Differences in terminal branch lengths could still cause some degree of negative correlations, and our method does not account for this. This may be a plausible explanation for some patterns: insecticide resistance has been known to spread across species in *Anopheles (Grau-Bové, Tomlinson et al. 2020)*, while sex chromosomes are thought to be unlikely to introgress (Fraïsse, Sachdeva 2021, Muralidhar, Coop 2024).

Overall, this study provides a new method which allows for the rapid identification of evolutionary rate correlations for a focal taxon, enabling the interrogation of a whole suite of questions. Our approach allows for the identification of gene-pairs whose evolutionary rates are correlated, identification of such pairs that exceed expectation under some degree of genome-wide correlations, and analyses of pathways and genes that are enriched for co-evolutionary signals for individual genes. We demonstrate the utility for a model system, and an extremely well-studied focal gene, but the method has been built with non-model systems and understudied genes in mind. As high-quality reference genomes for different taxa become increasingly numerous, this approach should be usable for any organism of interest.

## Acknowledgements

We’d like to thank the Dagilis lab for constructive feedback on the manuscript. AJD and BD were supported by start-up funds from the University of Connecticut. DRM was supported by R35GM148244.

## Data/script availability

All data used were publicly available reference genomes (Table S1). Scripts necessary to replicate this analysis are available in the GitHub repository (https://github.com/adagilis/CoEvolution) and a full guide to the pipeline can be found at (https://github.com/adagilis/CoEvolution/blob/main/scripts/Pipeline.pdf). Result files for analyses will be made available on figshare following peer review. Please reach out to Andrius Dagilis if you’d like any pre-review files.

## Supplementary Materials

Tables S1-S3,S5-S7 are available as Supplementary File 1.

**Table S3:** Genes with the highest fERC values.

| Protein (Flybase gene) | Mean fERC | Significant interactions | Name; Function |
| --- | --- | --- | --- |
| NP_001259732.1 (FBgn0031069) | 3.17 | 2265 | Abcd3; ATP binding cassette transporter. |
| NP_651601.1 (FBgn0039562) | 3.04 | 2313 | Gp93; a heat shock protein Hsp90 family member that is involved in midgut development. |
| NP_731967.2 (FBgn0266064) | 2.99 | 2243 | Glys; glycogen synthase |
| NP_524637.2 (FBgn0025741) | 2.98 | 2178 | PlexA; Plexin - alters actin, microtubule adhesion and axon development |
| NP_476610.2 (FBgn0002183) | 2.96 | 2080 | dre4; chromatin regulator, core factor in chromatin remodeling |
| NP_001286414.1 (FBgn0001220) | 2.95 | 2158 | Hsc70-5; Heat shock protein 70. |

**Figure S1:**
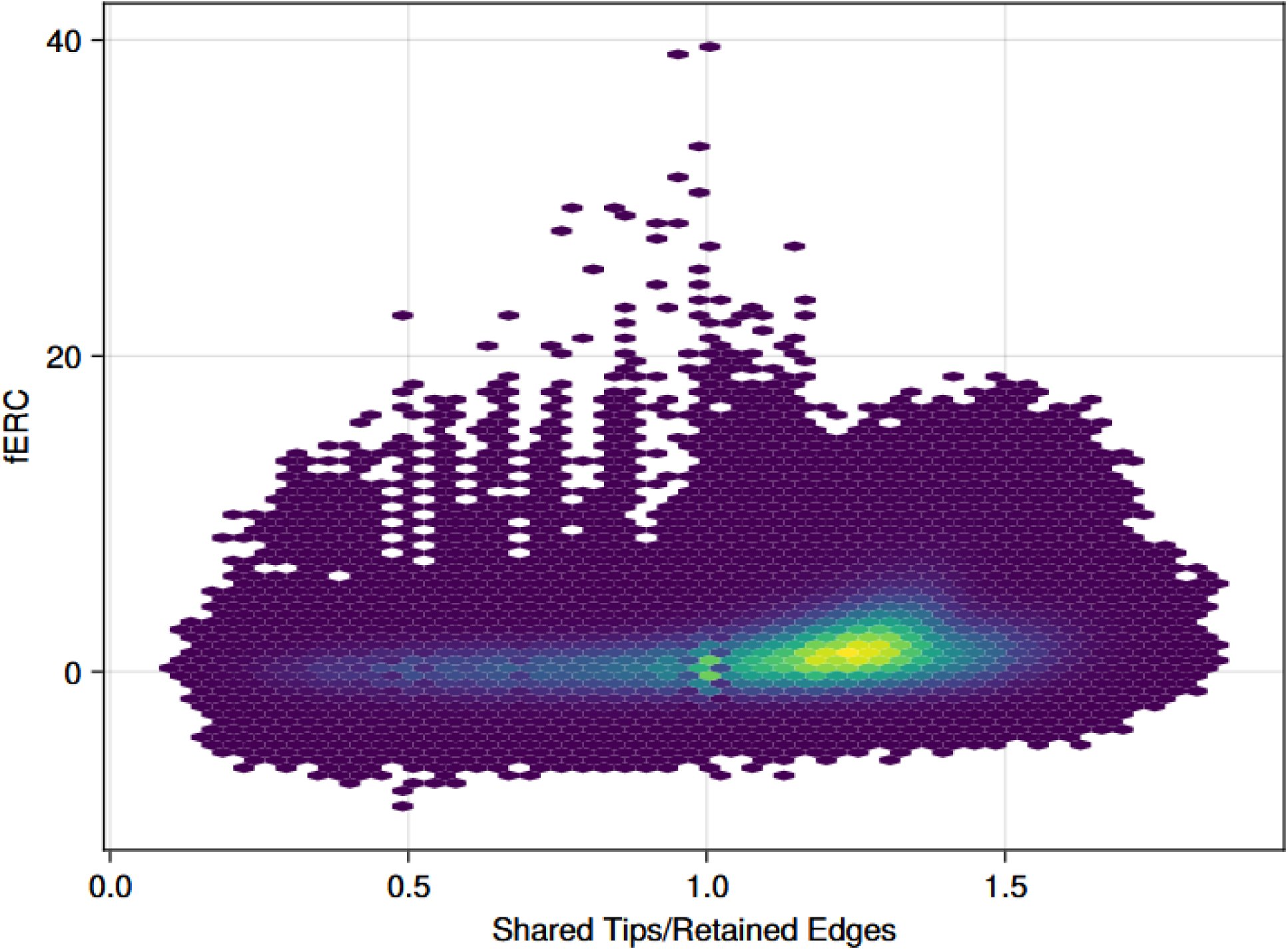
fERC increases as a function of the number of retained edges over the tips shared by a pair of trees. This suggests that trees that share topology, rather than simply many tips, are more likely to show evidence of co-evolution. Note that our most extreme values of fERC occur for pairs of trees in which the number of tips retained is very close or equal to the number of shared tips. These are all trees with very few tips in general, and therefore few possible edges.

**Figure S2:**
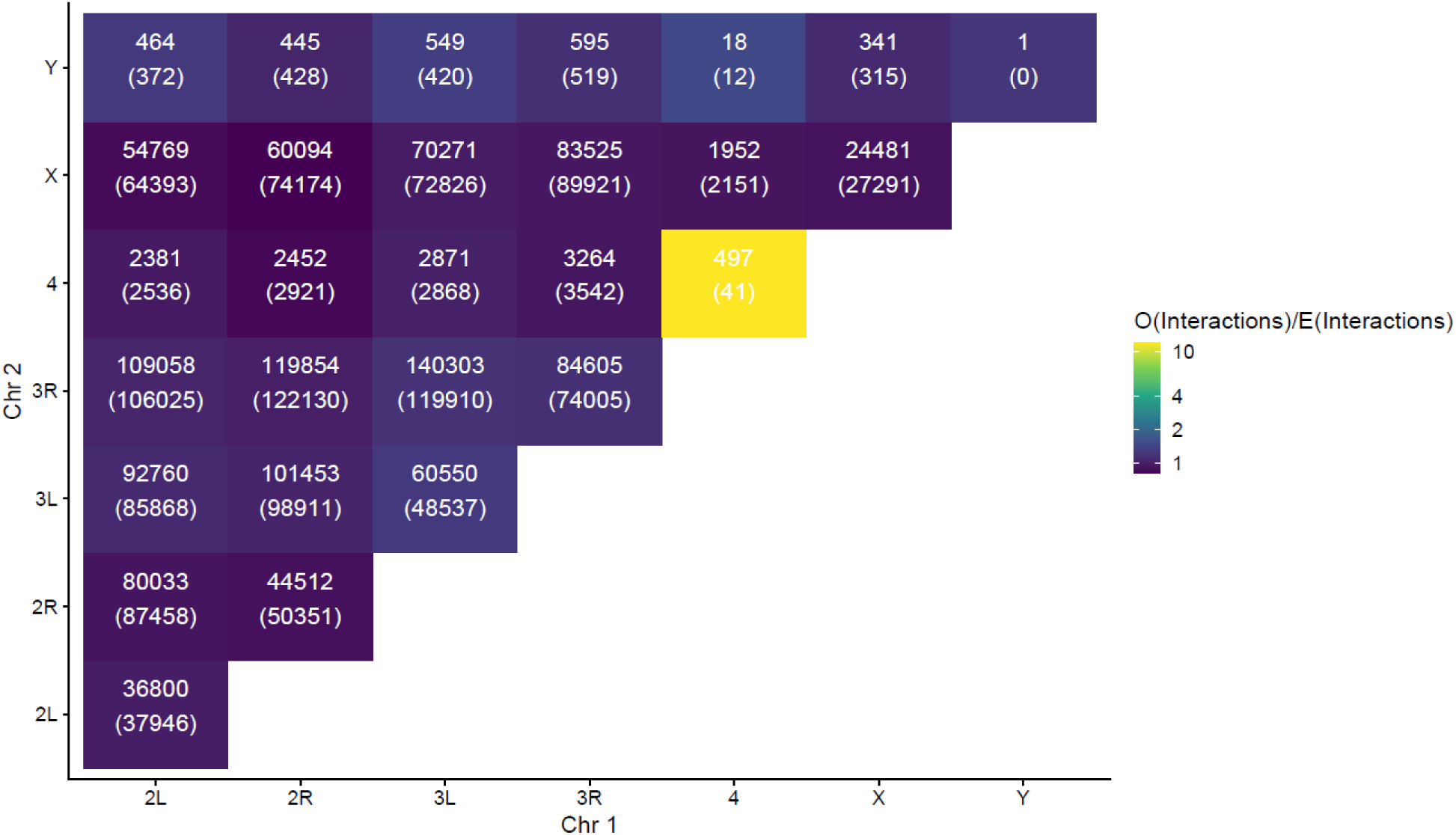
Observed vs Expected significant interactions between chromosomes. We calculate the expected number of interactions as the number of combinations of genes in our dataset between each pair of chromosomes multiplied by the fraction of total interactions that were significant. The ratio of observed to expected is indicated by the fill color, while the text shows the values of observed (top) and expected (bottom) pairs of genes with significant interactions. Overall, we note a lack of mitochondrial interactions, and an overrepresentation of interactions on Chromosome 4. However, in both cases, these are represented by relatively few genes, and so even a few genes having unusual numbers of orthologs/tree topology consistency can lead to overall decreases/increases in significant edges.

**Figure S3:**
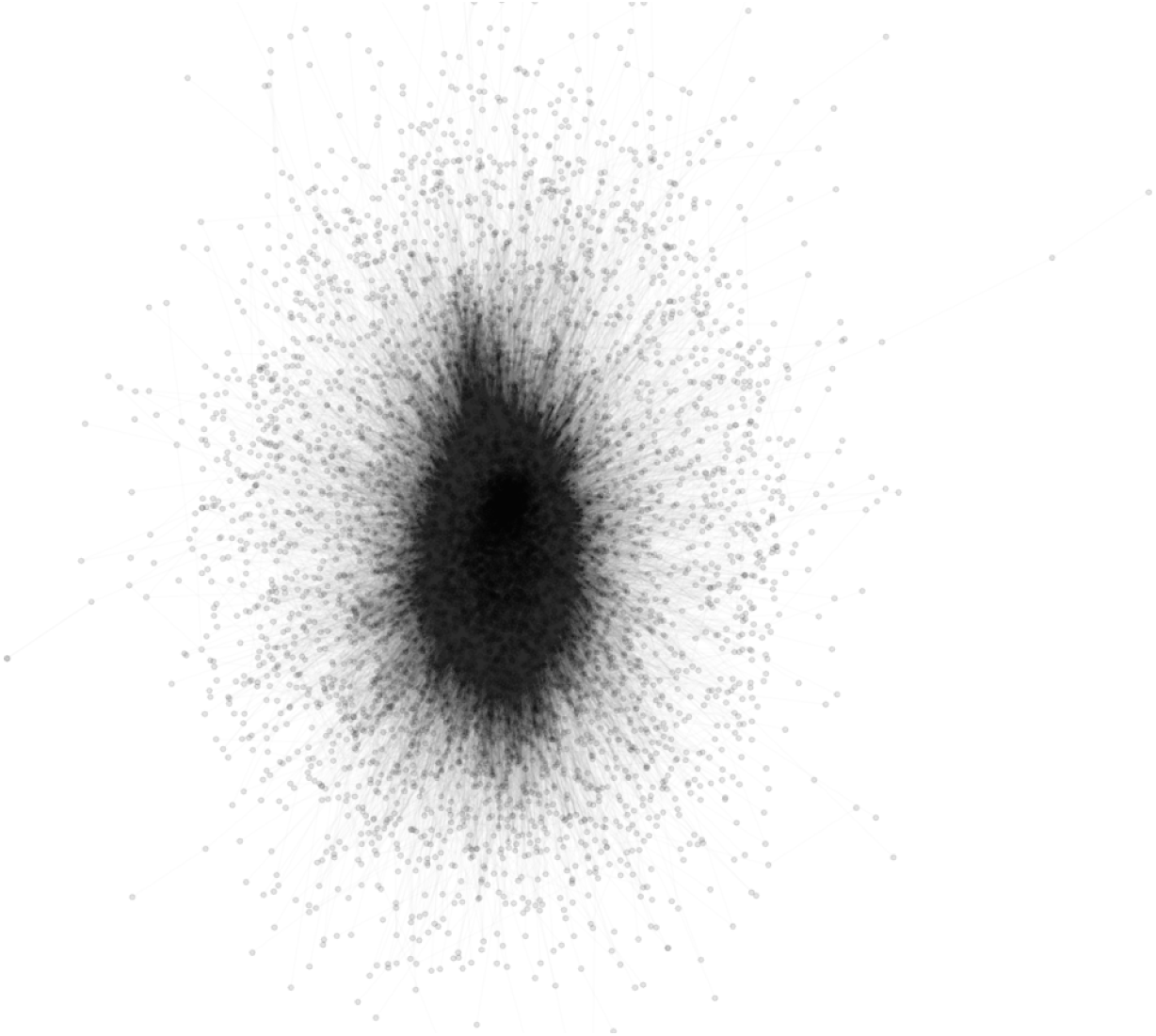
Network of significant co-evolutionary interactions. Each point represents an individual sequence, while lines are significant interactions. Only the largest connected component is shown for convenience.

**Figure S4:**
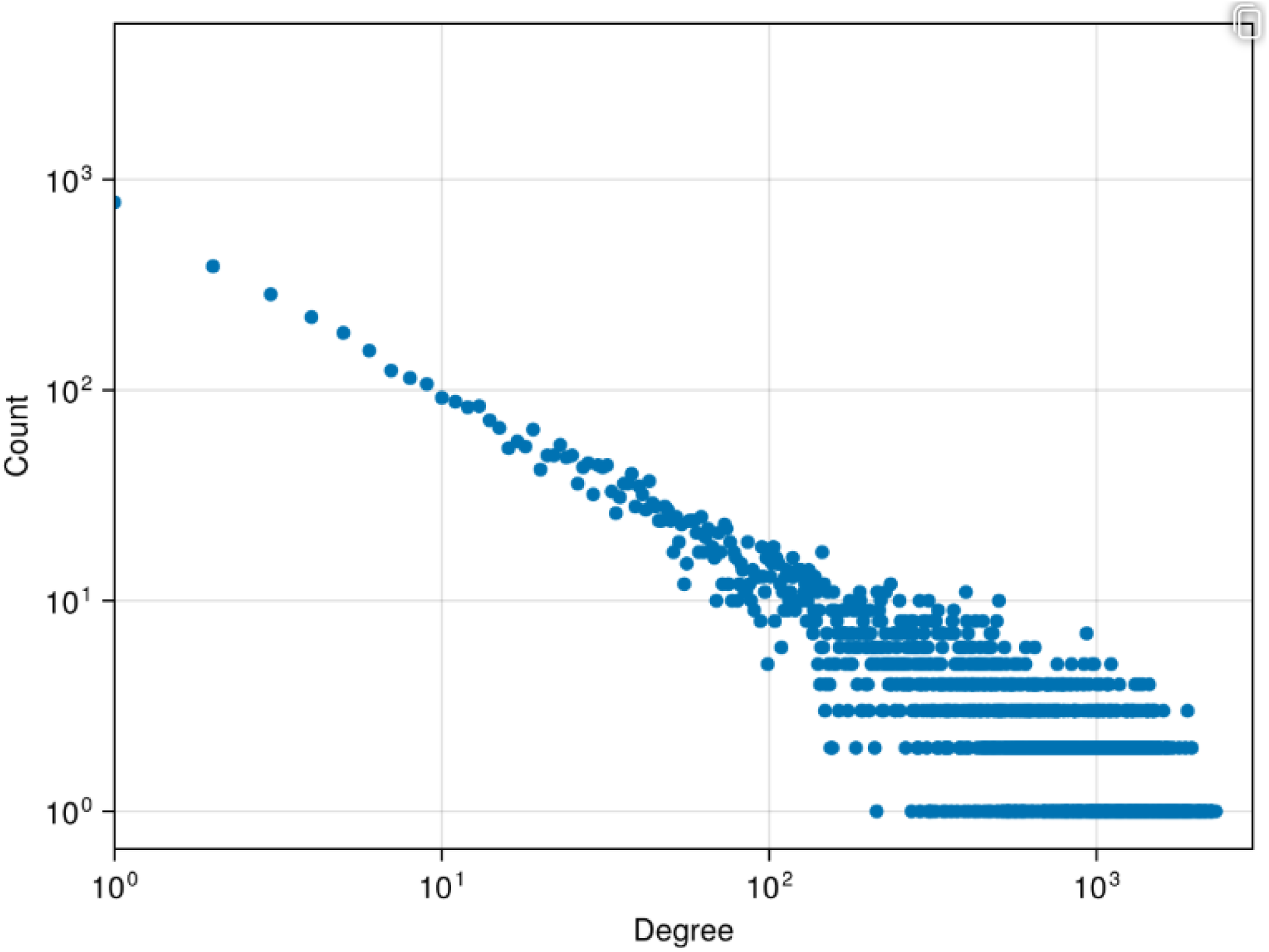
Degree distribution in network. The number of genes with degree (number of significant interaction partners) is distributed via Power Law, similar to other biological networks and opposed to networks in which interactions are distributed randomly (and therefore the degree distribution follows a binomial distribution).

**Figure S5:**
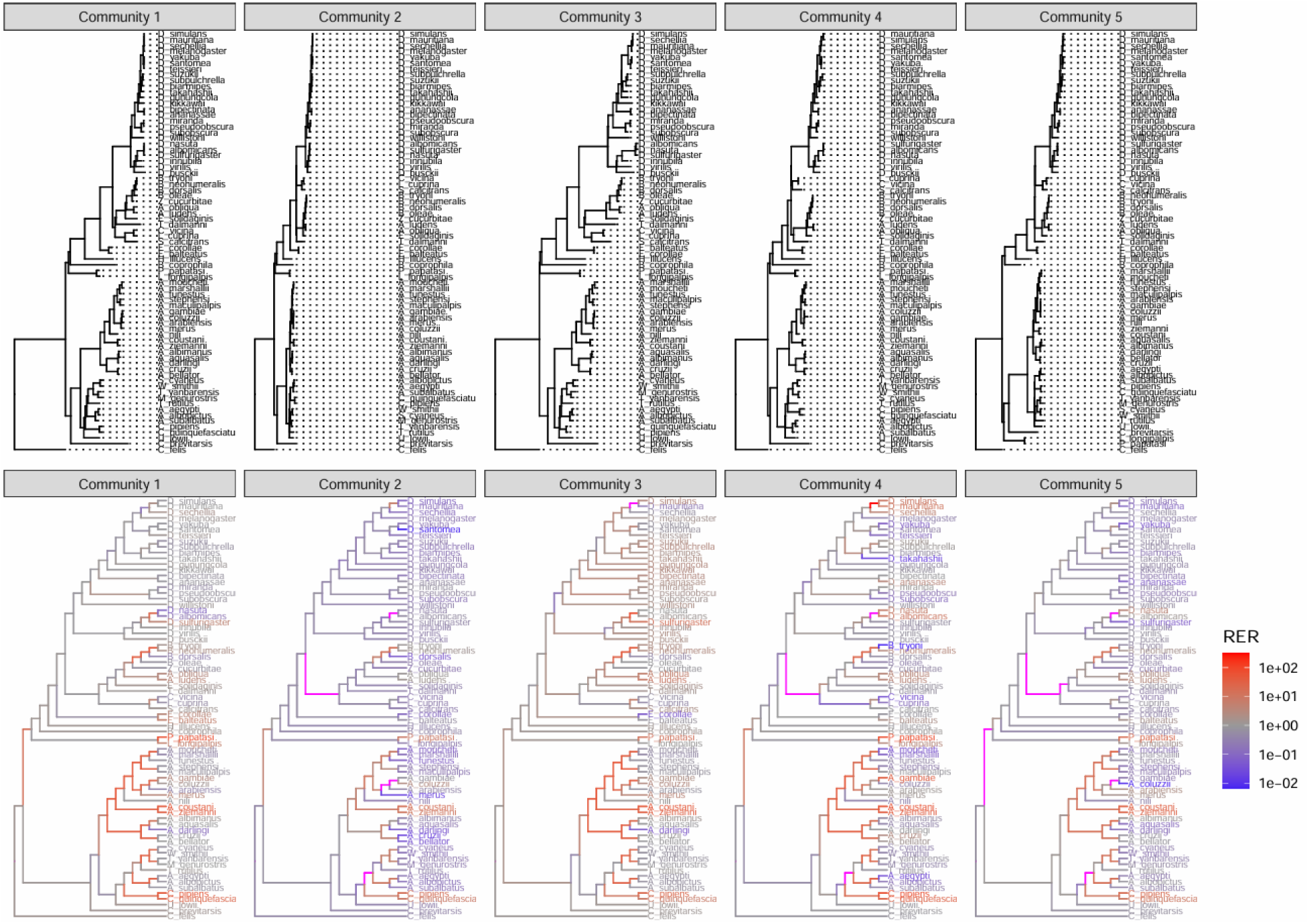
Consensus community phylogenies. For each of the clusters of genes with more than 50 members in our network, we generated an ASTRAL-IV consensus tree (A). To help visualize the individual branch rate differences/branches missing from the species tree, we plot the relative evolutionary rate for each branch (branch length in community tree divided by branch length in species tree) (B). Branches present in species tree but missing in gene tree are colored magenta. Note that Community 2 genes overall evolve more slowly than the species tree, while community 3 show rapid evolution in more recent tips.

**Figure S6:**
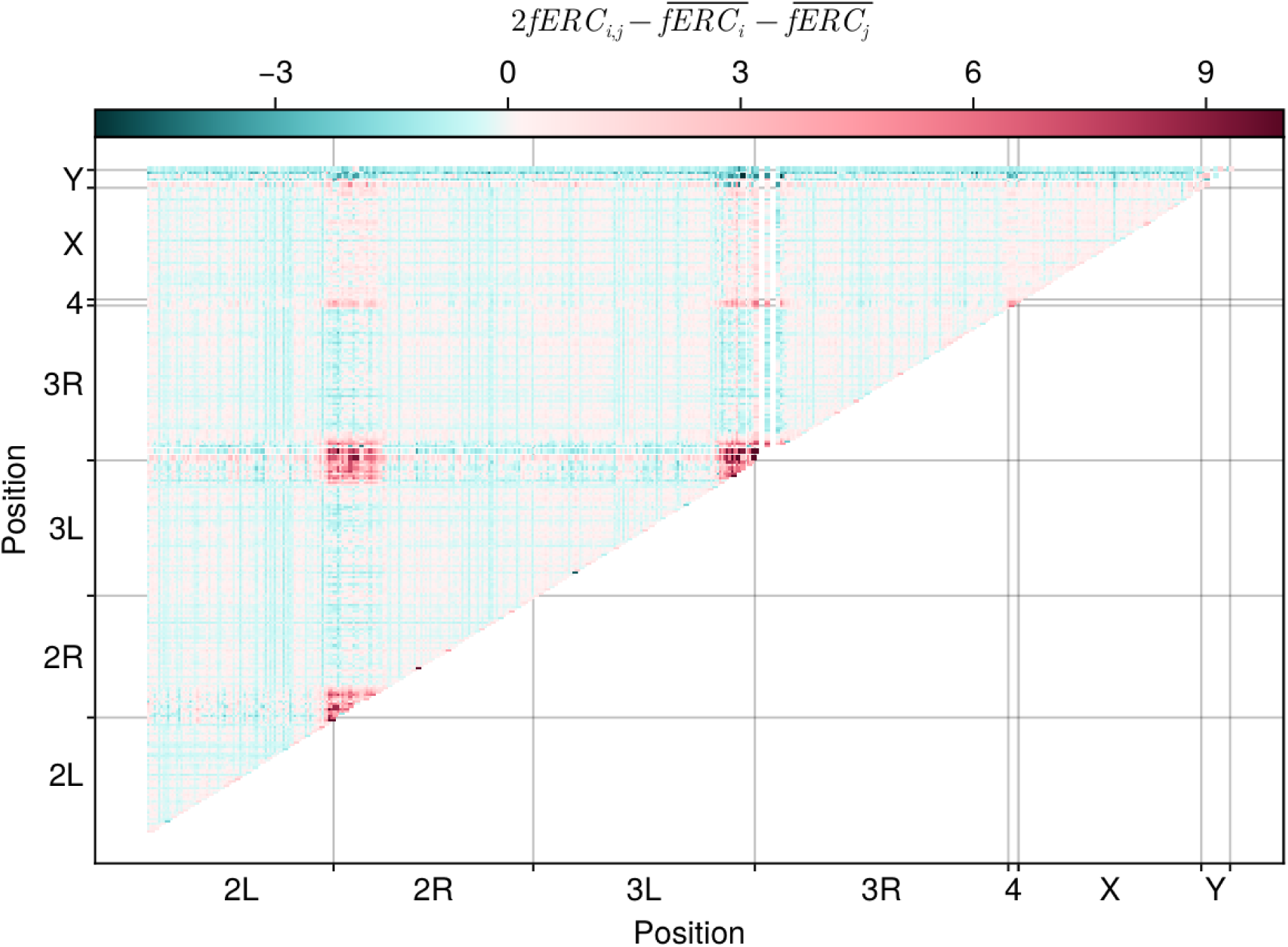
fERC specificity across the genome. For each interaction, we relativized its strength to the average interaction strength of both genes involved. The resulting values show how specific an interaction is compared to the average interaction of each gene. Note that the centromeres show highly specific interactions with each other (and chromosome 4), while most other interactions are either non-specific or nearly so. Interactions with the Y, in general, seem to be weaker than expected given the rest of the interaction network.

